## Supplemental Information for "Confinement of unliganded EGFR by tetraspanin nanodomains gates EGFR ligand binding and signaling"

Sugiyama et al.

### Supplemental Information

Figure S1-S12

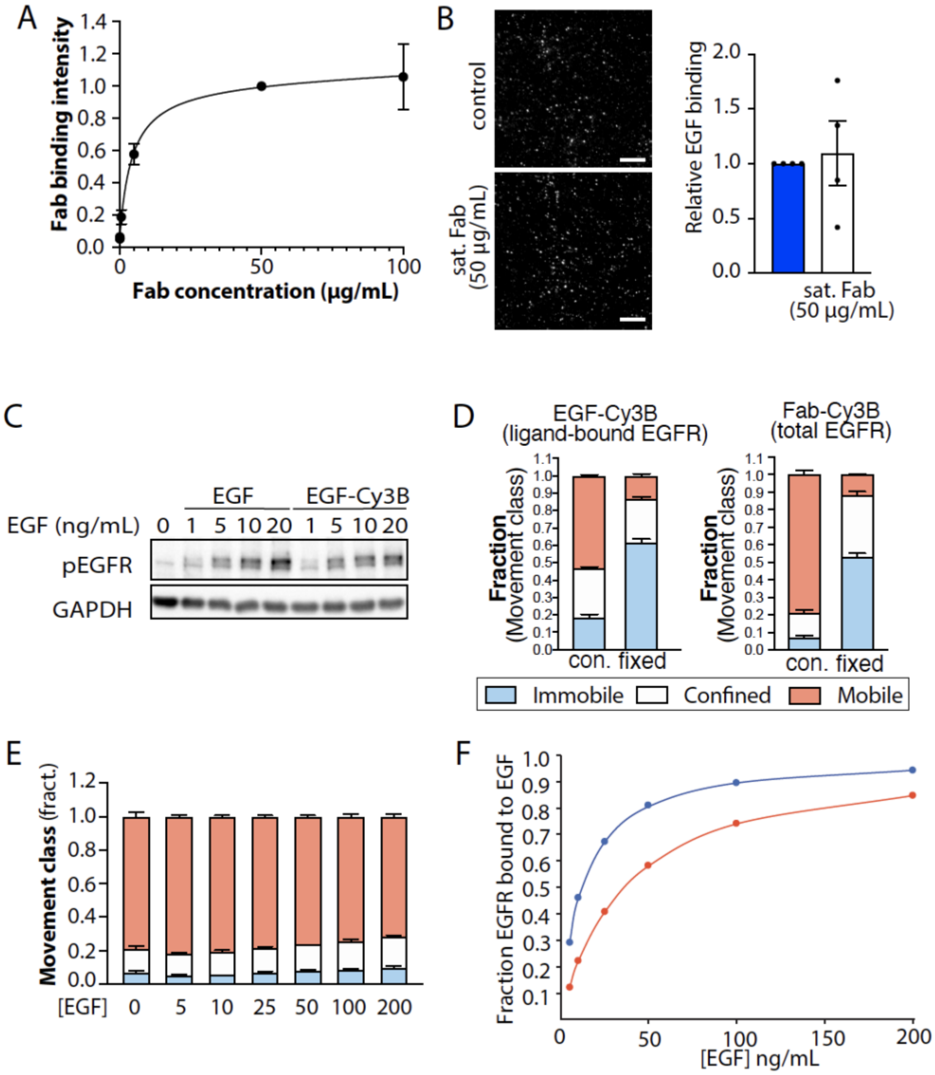


**Figure S1. Validation of labeling and tracking strategies used for single-particle tracking**. (A) Intact ARPE-19 cells were treated with Fab-Cy3B at various concentrations, followed by fixation, imaging using widefield epifluorescence microscopy, and measurement of cell-associated Fab-Cy3B; shown are the means ± SE. (B) ARPE-19 cells were treated with Fab-Cy3B under saturating conditions (50 µg/mL), or left untreated, prior to labeling for 2 min with Cy3B-EGF. Shown are representative images as well as the quantification of cell surface EGF-Cy3B from n=4 independent experiments, showing the overall mean (bar) ± SE, and mean values from independent experiments (dots). Scale = 20 µm (C) ARPE-19 cells were stimulated with indicated concentrations of EGF-Cy3B or unlabelled EGF for 5 min. Shown are immunoblots of whole-cell lysates detecting phosphorylated EGFR (pY1068) or GAPDH (loading control). (D) ARPE-19 cells were subjected to methanol fixation (fixed) or not (control, con.), followed by labeling with either EGF-Cy3B or Fab-Cy3B. Results of SPT analysis showing the fraction of all EGFR tracks, as labelled by Fab-Cy3B (left panels), or showing the fraction of only ligand-bound EGFR, as labeling EGF-Cy3B (right panels) that exhibit mobile, confined or immobile behaviour. (E) Extended results of SPT analysis as per Fig. 1. Shown are the mean ± SE of the fraction of all EGFR tracks, as labelled by Fab- that exhibit mobile, confined or immobile behaviour. (F) Calculation of the fraction of EGFR bound to EGF at different concentrations based on a Kd of 2 nM (red line) or 6 nM (blue line). This is based on Michaelis-Menten equation, which provides that Fraction bound = (Bmax)[L]/([L]+Kd), and the assumption that Bmax = 1.

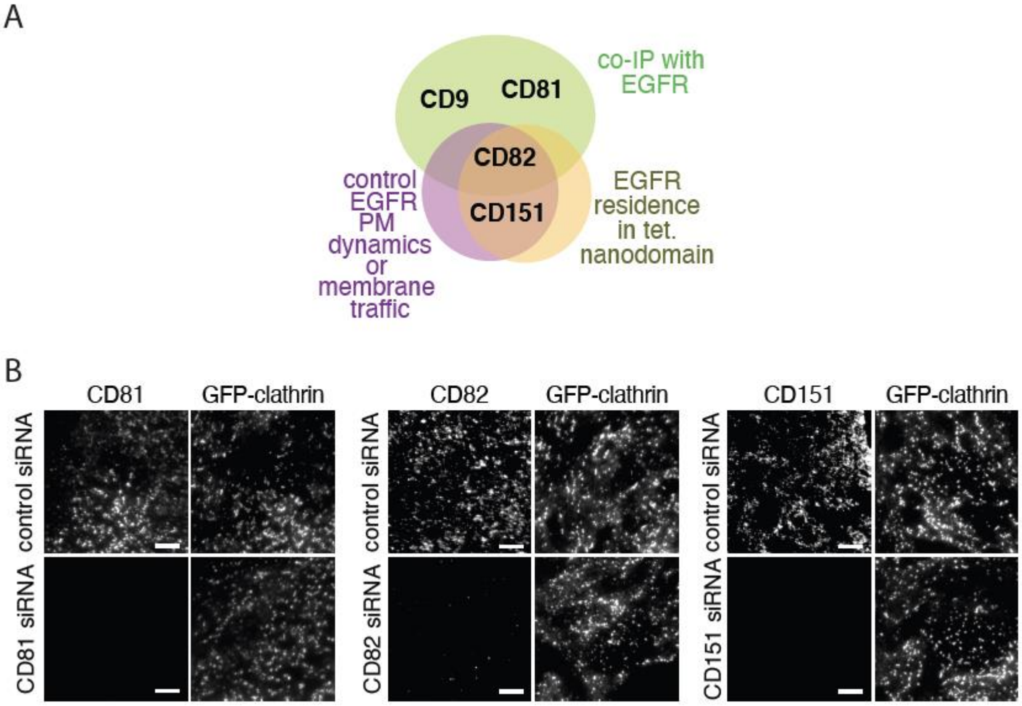


**Figure S2. Tetraspanin interactions with EGFR and tetraspanin antibody validation.** (A) Diagram showing the previously published interactions (physical or functional) of 4 tetraspanins (CD9, CD81, CD82 and CD151) with EGFR. (B) Antibody validation for detection of CD151 or CD82 via TIRF microscopy. ARPE-19 cells stably expressing GFP-clathrin were treated with siRNA targeting CD81, CD82 or CD151, followed by labeling with either anti-CD81, anti-CD82 or -CD151 antibodies, as indicated. Shown are representative images obtained by TIRF-M. scale 5 μm.

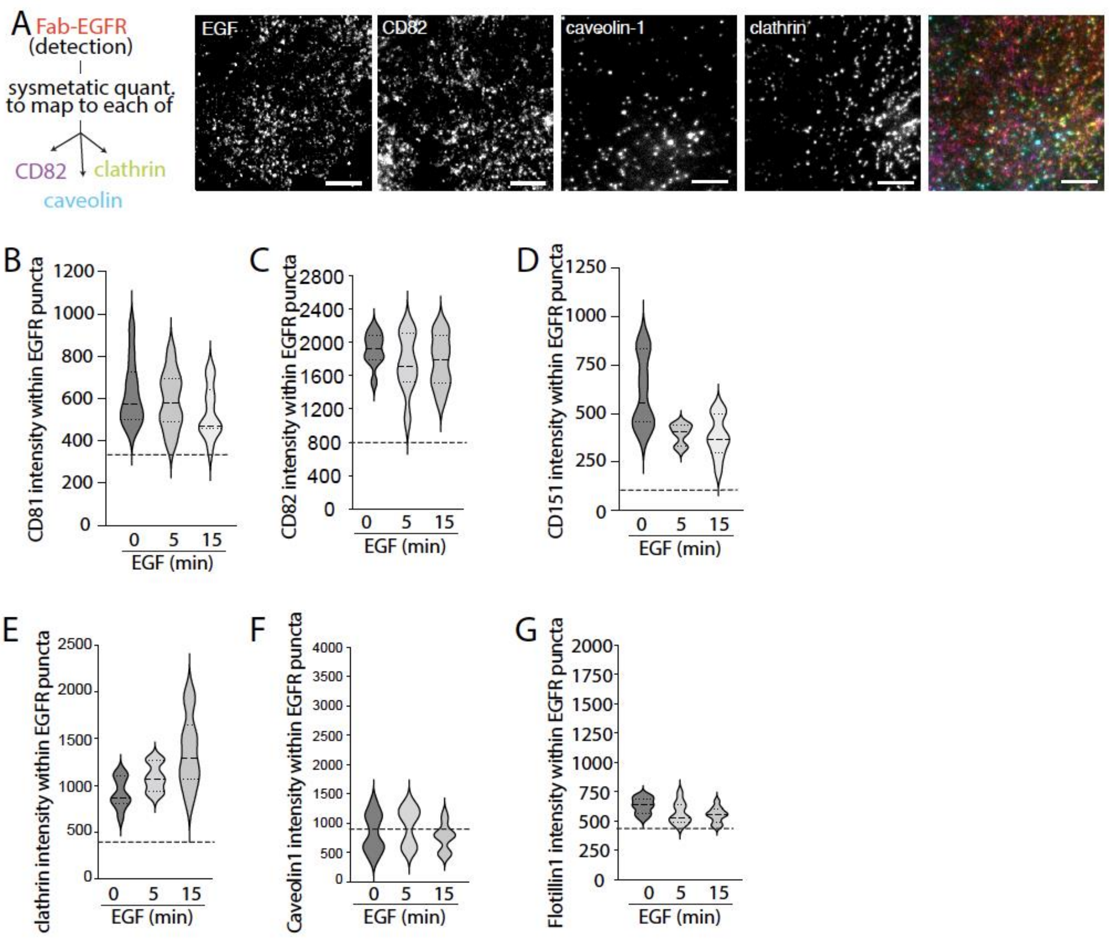


**Figure S3. Labeling of multiple nanodomain markers alongside EGFR**. (A) Multi-channel parallel labeling of EGFR with multiple nanodomain markers. Diagram showing labeling strategy (left panels), which involves labeling of cells with Fab-Cy3B (to label total EGFR), followed by fixation and staining nanodomain markers. Shown are representative images obtained by TIRF-M of ARPE-19 stably expressing eGFP-clathrin labeled by IF staining for CD82 and caveolin-1, followed by labeling with Cy3B-Fab (to detect total EGFR). (B-G) Shown are results of detection of EGFR objects followed by *intensity-based analysis* of EGFR object overlap with the indicated secondary channel (tetraspanins, clathrin, flotillin1, caveolin1) (as described in Methods). Also shown is the background overlap (horizonal dashed line) determined by repeating measurements of EGFR overlap with each secondary marker following rotation of one image by 180 degrees to randomize the marker overlap. Results are shown as the distribution of measurements in individual cells (violin plot), featuring the 25$^{th}$, 50$^{th}$ and 75$^{th}$ percentiles (horizonal dashed lines).

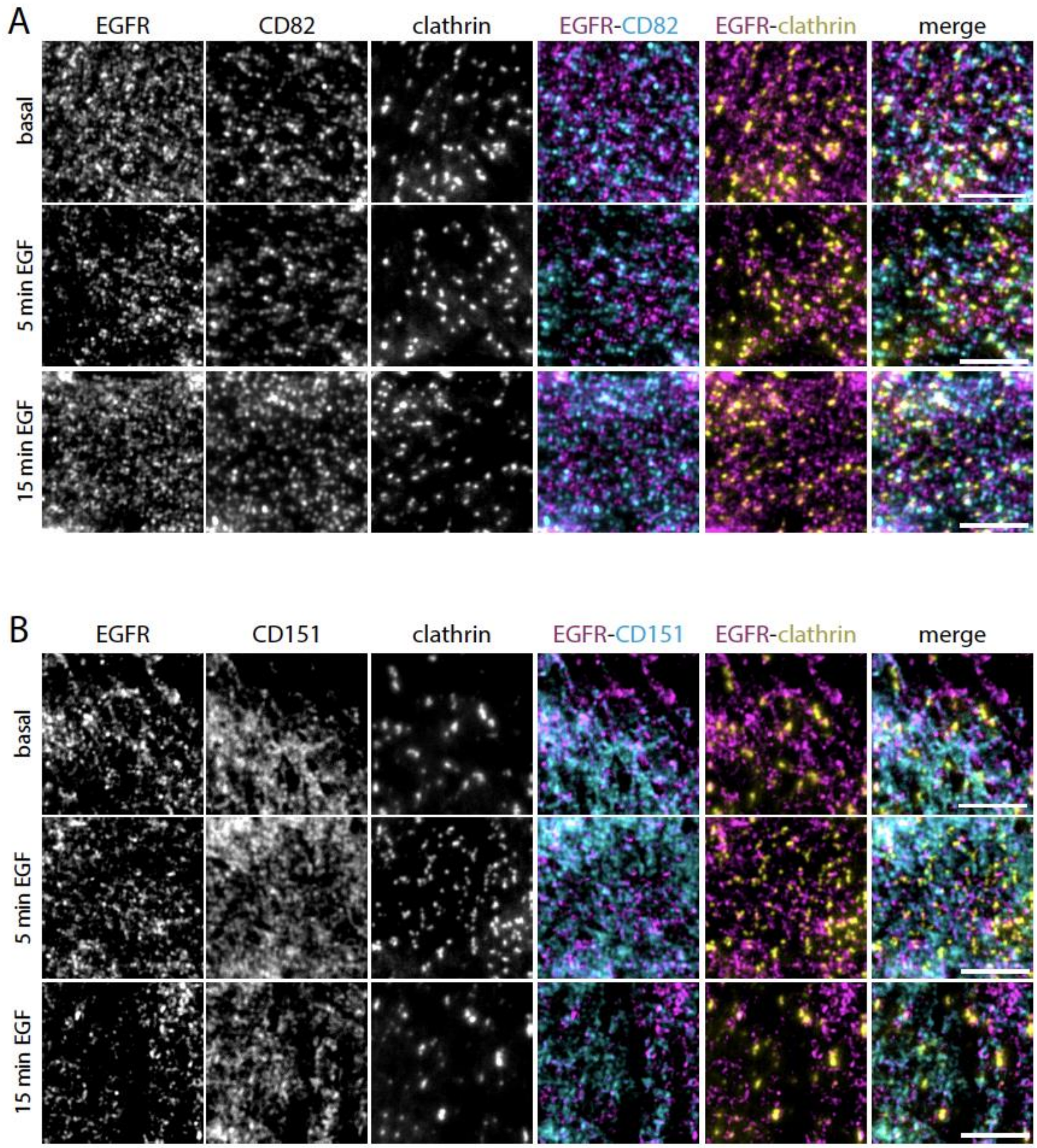


**Figure S4. Localization of EGFR relative to CD82, CD151 and clathrin at the cell surface**. ARPE-19 cells stably expressing eGFP-clathrin were labeled with Fab-Cy3B (to label total EGFR) and stimulated with EGF as indicated, followed by fixation and staining with CD82 (A) or CD151 (B) antibodies. Shown for each are representative images obtained by TIRF-M; antibody labeling of tetraspanins is highly specific (**Fig. S2**); similar experiments with labeling of CD81 were performed (**Fig. 2**). Scale = 5 μm.

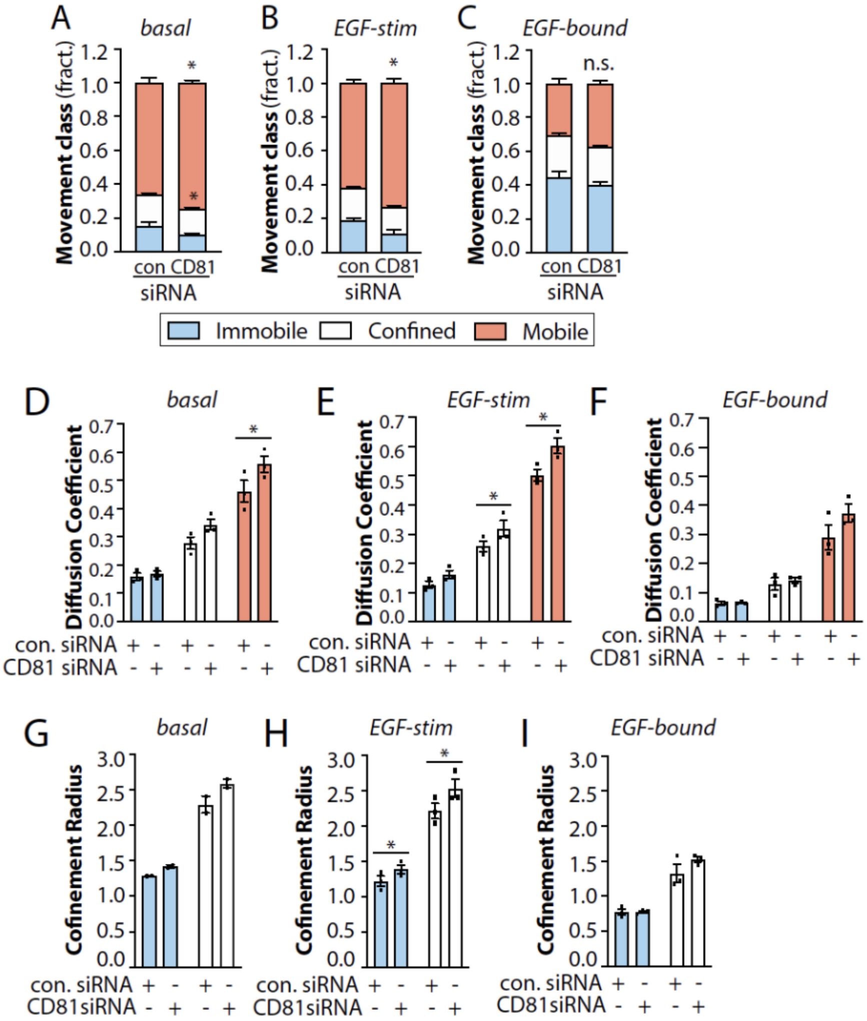


**Figure S5. Silencing CD81 impacts the diffusion coefficient of mobile EGFR in MDA-MB-231 cells.** MDA-MB-231 cells were treated with siRNA to silence CD81 or non-targeting siRNA (control), as indicated. (A-I) Results of SPT analysis. The cells were then subjected to SPT using either Fab-Cy3B to label total EGFR in the absence (A, D, G) or presence (B, E, H) of unlabelled EGF, or labelled using EGF-Cy3B (C, F, I), to label only ligand-bound EGFR). Shown in A-C are the mean ± SE of the fraction of EGFR tracks in each mobility category (immobile, confined, mobile) under each condition, as well as in A-B the same data re-plotted to view the immobile and confined fractions. Also shown are mean ± SE of diffusion coefficient (D-F) or the confinement radius (G-I). EGF-Cy3B (ligand-bound) data is from 3 independent experiments, and Fab-Cy3B (total EGFR) data is from 5 independent experiments. Each experiment involved detection and tracking of >500 EGFR objects. *, $p < 0.05$

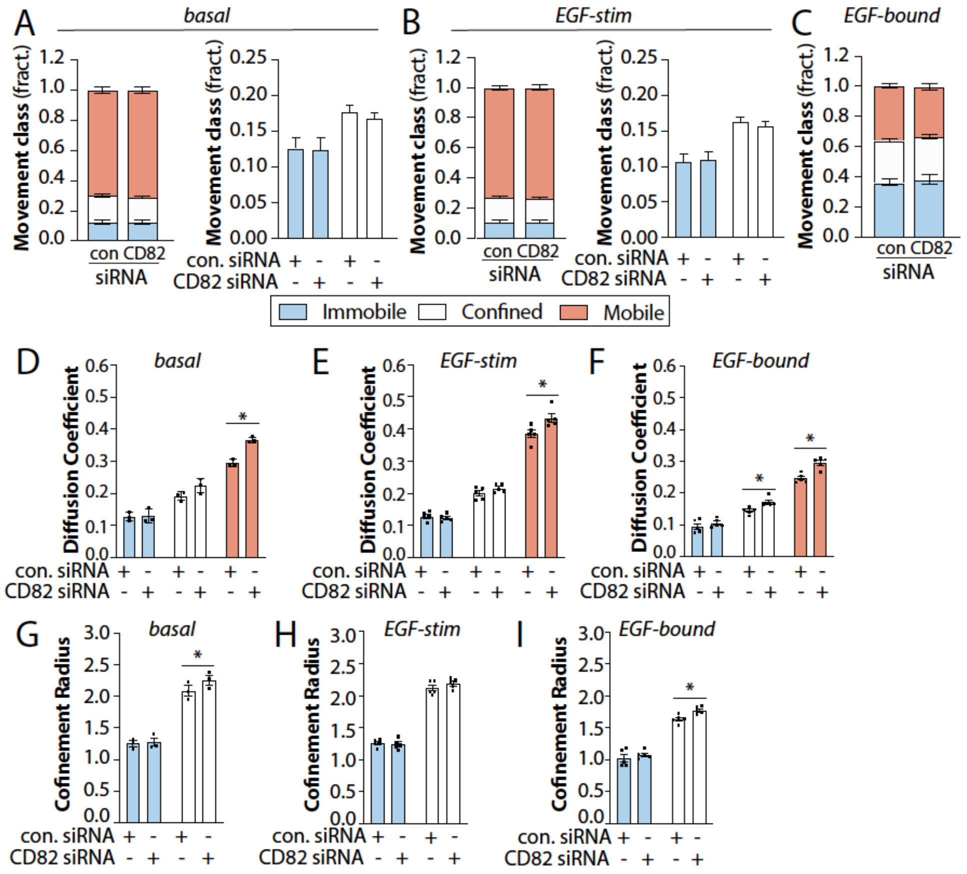


**Figure S6. Silencing CD82 impacts the diffusion coefficient of mobile EGFR.** ARPE-19 cells were treated with siRNA to silence CD82 or non-targeting siRNA (control), as indicated. (A-I) Results of SPT analysis. The cells were then subjected to SPT using either Fab-Cy3B to label total EGFR in the absence (A, D, G) or presence (B, E, H) of unlabelled EGF, or labelled using EGF-Cy3B (C, F, I), to label only ligand-bound EGFR). Shown in A-C are the mean ± SE of the fraction of EGFR tracks in each mobility category (immobile, confined, mobile) under each condition, as well as in A-B the same data re-plotted to view the immobile and confined fractions. Also shown are mean ± SE of diffusion coefficient (D-F) or the confinement radius (G-I). EGF-Cy3B (ligand-bound) data is from 3 independent experiments, and Fab-Cy3B (total EGFR) data is from 5 independent experiments. Each experiment involved detection and tracking of >500 EGFR objects. *, $p < 0.05$

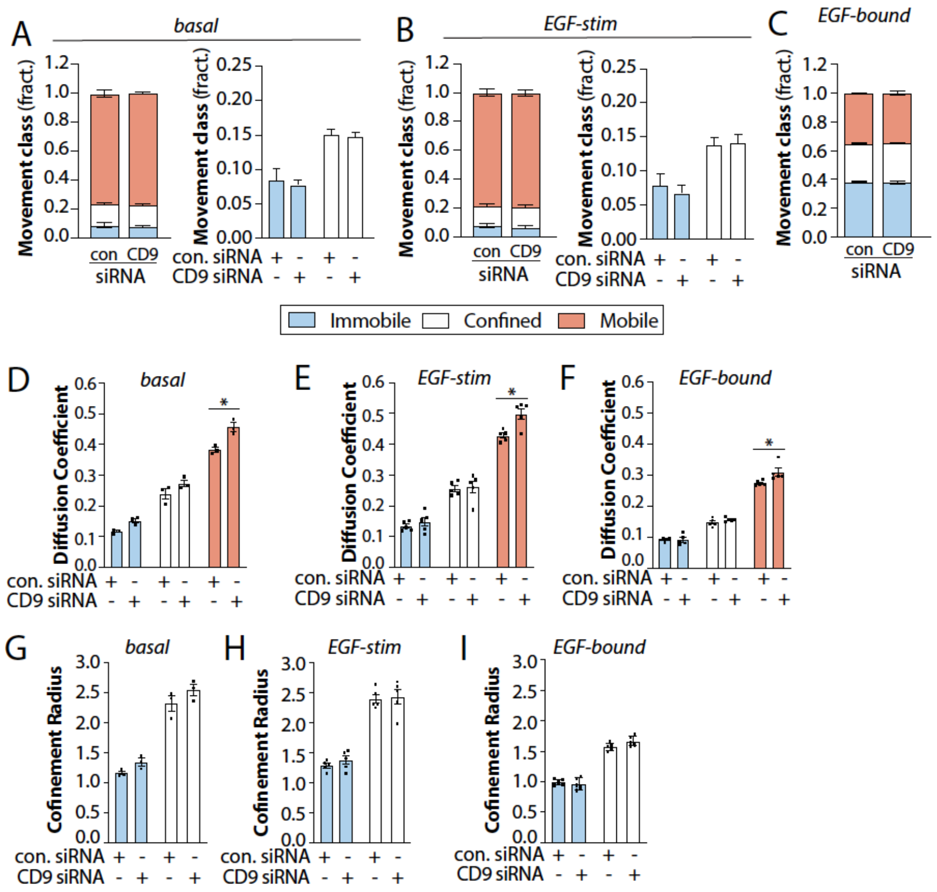


**Figure S7. Silencing CD9 impacts the diffusion coefficient of mobile EGFR.** ARPE-19 cells were treated with siRNA to silence CD9 or non-targeting siRNA (control), as indicated. (A-I) Results of SPT analysis. The cells were then subjected to SPT using either Fab-Cy3B to label total EGFR in the absence (A, D, G) or presence (B, E, H) of unlabelled EGF, or labelled using EGF-Cy3B (C, F, I), to label only ligand-bound EGFR). Shown in A-C are the mean ± SE of the fraction of EGFR tracks in each mobility category (immobile, confined, mobile) under each condition, as well as in A-B the same data re-plotted to view the immobile and confined fractions. Also shown are mean ± SE of diffusion coefficient (D-F) or the confinement radius (G-I). EGF-Cy3B (ligand-bound) data is from 5 independent experiments, and Fab-Cy3B (total EGFR) data is from 3 independent experiments. Each experiment involved detection and tracking of >500 EGFR objects. *, $p < 0.05$

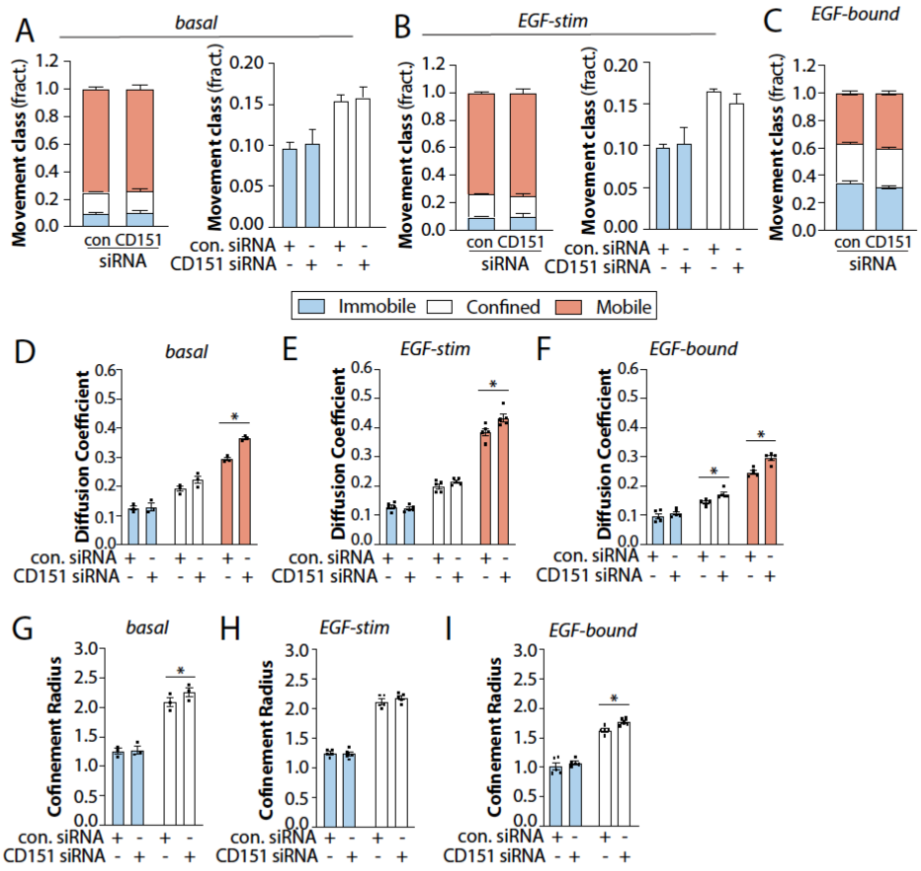


**Figure S8. Silencing CD151 impacts the diffusion coefficient of mobile EGFR.** ARPE-19 cells were treated with siRNA to silence CD151 or non-targeting siRNA (control), as indicated. (A-I) Results of SPT analysis. The cells were then subjected to SPT using either Fab-Cy3B to label total EGFR in the absence (A, D, G) or presence (B, E, H) of unlabelled EGF, or labelled using EGF-Cy3B (C, F, I), to label only ligand-bound EGFR). Shown in A-C are the mean ± SE of the fraction of EGFR tracks in each mobility category (immobile, confined, mobile) under each condition, as well as in A-B the same data re-plotted to view the immobile and confined fractions. Also shown are mean ± SE of diffusion coefficient (D-F) or the confinement radius (G-I). EGF-Cy3B (ligand-bound) data is from 5 independent experiments, and Fab-Cy3B (total EGFR) data is from 3 independent experiments. Each experiment involved detection and tracking of >500 EGFR objects. *, $p < 0.05$

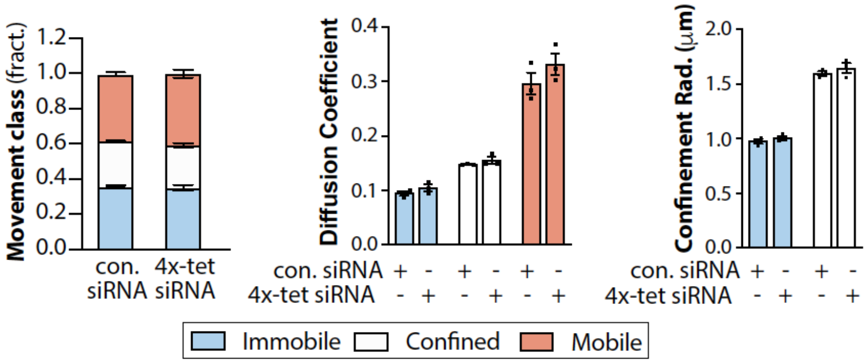


**Figure S9. Concomitant silencing of multiple tetraspanins does not alter mobility of ligand-bound EGFR**. ARPE-19 cells were treated with siRNA to silence individual tetraspanin proteins, or CD9, CD81, CD82 and CD151 concomitantly (4x-tet), or non-targeting siRNA (control). The cells were then subjected to SPT using EGF-Cy3B to label ligand-bound EGFR. Shown are the results of the SPT analysis, showing the fraction of EGFR tracks in each mobility category (immobile, confined, mobile) under each condition (left panel), diffusion coefficient of EGFR by mobility class (middle panel) and confinement radius of EGFR in immobile and confined EGFR populations (right panel).

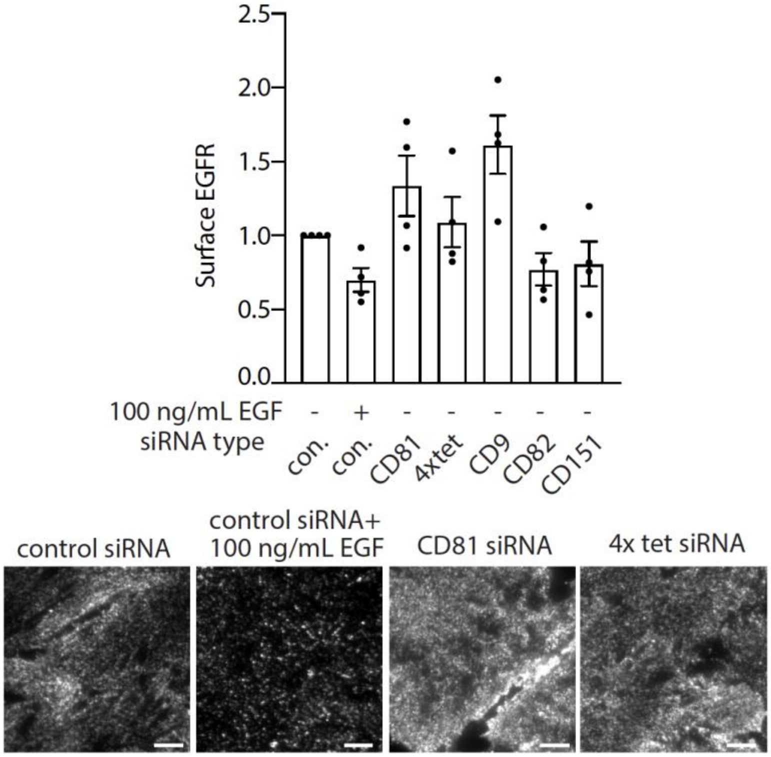


**Figure S10. Measurement of cell surface EGFR levels following tetraspanin silencing**. ARPE-19 cells were transfected with siRNAs targeting tetraspanins as indicated. Following transfection, intact (non-permeabilized) cells were labelled with mAb108 (detecting surface-exposed EGFR), followed by fixation and labeling with appropriate secondary antibodies. Shown are representative microscopy images (bottom panels) as well as measurement of cell surface EGFR labeling intensity, showing the overall mean (bar) ± SE, as well as the mean from individual experiments (dots). Scale = 20 μm.

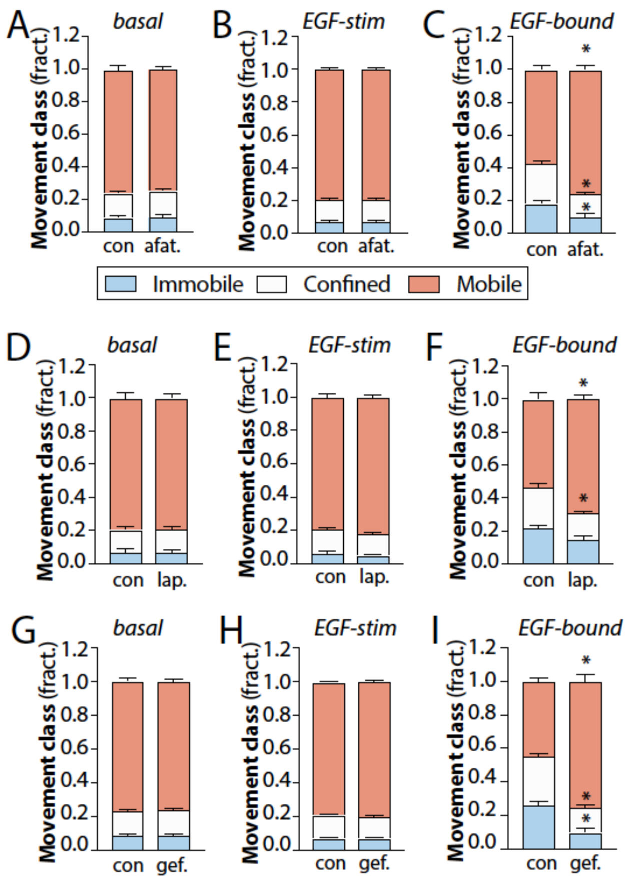


**Figure S11. EGFR tyrosine kinase inhibitors (TKIs) selectively impact confinement of ligand-bound EGFR**. ARPE cells were treated with 2 µM of each TKI as indicated. The cells were then subjected to SPT using either Fab-Cy3B to label total EGFR in the absence (A, D, G) or presence (B, E, H) of unlabelled EGF, or labelled using EGF-Cy3B (C, F, I), to label only ligand-bound EGFR). Shown are the mean ± SE of the fraction of EGFR tracks in each mobility category (immobile, confined, mobile) under each condition. All data is from 3 independent experiments. Each experiment involved detection and tracking of >500 EGFR objects. *, $p < 0.05$

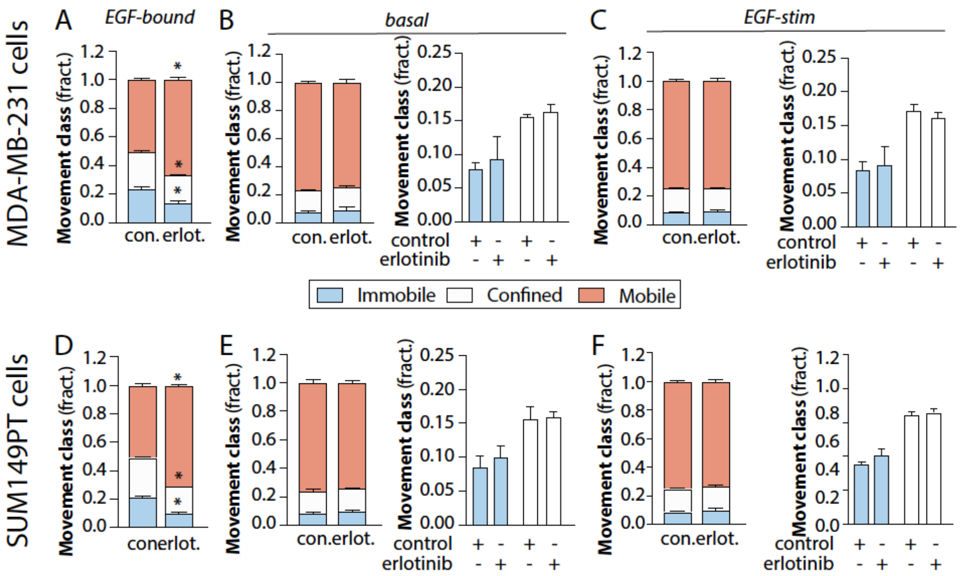


**Figure S12. Erlotinib selectively impacts mobility of ligand-bound EGFR in MDA-MB-231 and SUM149PT breast cancer cells**. MDA-MB-231 cells (A-C) or SUM149-PT cells (D-F) were pre-treated with 2 µM erlotinib for 20 min. The cells were then subjected to SPT using either using EGF-Cy3B (A,B), to label only ligand-bound EGFR) or Fab-Cy3B to label total EGFR in the absence (B, E) or presence (C, F) of unlabelled EGF, or. Shown are the mean ± SE of the fraction of EGFR tracks in each mobility category (immobile, confined, mobile) under each condition, as well as in B-C, E-F the same data re-plotted to view the immobile and confined fractions. All data is from 3 independent experiments. Each experiment involved detection and tracking of >500 EGFR objects. *, $p < 0.05$
